# Towards Real-Time Airborne Pathogen Sensing: Electrostatic Capture and On-Chip LAMP Based Detection of Airborne Viral Pathogens

**DOI:** 10.1101/2024.01.05.574431

**Authors:** Nitin Jayakumar, Veronique Caffrey, Michael Caffrey, Igor Paprotny

## Abstract

Considerable loss of life, economic slowdown, and public health risk associated with the transmission of airborne respiratory pathogens was underscored by the recent COVID-19 pandemic. Airborne transmission of zoonotic diseases such as the highly pathogenic avian influenza (HPAI) and porcine reproductive and respiratory syndrome virus (PRRSV) has caused major disruptions to domestic and global food security. Current ambient air pathogen monitoring systems involves the collection of air samples from indoor settings suspected of viral contamination, followed by subsequent processing of capture samples to determine the presence and species of airborne viral matter. Nucleic acid amplification techniques are considered the gold standard for pathogen diagnostics. Currently, the necessary extraction and purification of viral RNA from air collector systems prior to sample analysis is both time consuming and performed manually. A monitoring system with separate air sampling and biochemical detection procedures is prone to delay the response to emergent viral threats. In this paper, we present a pathogen monitoring system that overcomes these limitations related to extraction and purification of viral samples and lays the groundwork for a real-time monitor for airborne viral pathogens. We demonstrate a high flow electrostatic precipitator system, that uses small collection wells as counter electrodes for pathogen collection. Integrated reverse-transcriptase loop-mediated isothermal amplification (RT-LAMP) is used for detection of captured viral matter within wells. On-chip heating of collection wells is enabled by integrated planar heaters and small volumes of reagent (30 μ*L*) directly to the collection wells. We present the design of such a system and show experimental results that demonstrate the use of this device for detection of aerosolized SARS-CoV-2 virus like particles (VLPs), a model pathogen for SARV-CoV-2.

## 1. Introduction

COVID-19 pandemic demonstrated the considerable public health threat posed by airborne transmission of viruses. Besides the dissemination of the SARS-CoV-2 virus [1], various airborne pathogens, including influenza viruses, rhinovirus, measles virus, RSV, SARS-CoV, and MERS-CoV, have been identified in breath samples collected in indoor environments [2-7]. In the wake of the COVID-19 pandemic, numerous epidemiological studies have investigated aerosol suspension in diverse indoor settings such as restaurants [8,9], buses [10], cruise ships [11], dental clinics [12], and places of worship [13]. The collective findings of these studies implicate airborne aerosols as the primary transmission mode for a range of respiratory pathogens in human congregate settings. Furthermore, in animal settings, the airborne transmission of highly pathogenic avian influenza (HPAI) within the poultry farms and porcine reproductive and respiratory syndrome virus (PRRSV) in swine farms is also of great concern due to the high economic burden associated with outbreak control, disease eradication, and overall risk to food security. Between 2005 and 2021, HPAI has caused the mass depopulation of approximately 300 million poultry across 50 countries [14]. The 2015 HPAI outbreak in the US affected 232 farms in 15 states in less than 7 months, infecting 50 million birds and resulting in economic losses valued at $3.3 billion [15]. HPAI transmission in poultry farms have been attributed primarily to <u>airborne</u> particulate matter particles generated from feed, litter, feces, and animal feathers [15-18]. In-field epidemiological studies monitoring swine farms suggest airborne aerosols generated from the breathing, coughing, and sneezing of infected pigs as a possible transmission route for the spread of porcine reproductive and respiratory syndrome virus (PRRSV), in particular over distances less than 3 km [38,39].

### 1.1. Need for Real-Time Airborne Viral Monitoring of Indoor Settings

Viral surveillance methods during the COVID-19 pandemic were predominantly conducted using nasal swab, and saliva-based point-of-care antigen tests. Serological data has since estimated that for every diagnosed case, there were 4.8 undiagnosed cases of SARS-CoV-2 infections [19], suggesting that these methods were crude, inaccurate, and difficult to implement. Similarly, tracheal, or cloacal swabs-based testing is currently the recognized standard for the monitoring of HPAI transmission in bird populations. In the case of avian populations, along with being time consuming, labor intensive, and difficult to scale, swab sampling is ineffective during early stages, when only a few animals are sick [16].

To address these monitoring challenges, airborne sampling in indoor settings has been adopted as an alternative approach towards viral surveillance and has been demonstrated for airborne collection and detection of SARS-CoV-2, influenza, and HPAI, and PRRSV viruses [20,21,38,43]. In these studies, viral RNA samples are captured through air filtration. Subsequently, filters are processed utilizing nucleic acid amplification techniques (NAA) for viral detection. Ramuta.et al [21] demonstrated the utility of filter-based air sampling and NAA based detection in real-world indoor settings (schools, hospitals, coffee shops, etc.). Additionally, in the same study, genomic sequencing of air samples confirmed the ability to track emerging variants, by frequent sampling of indoor air utilizing commercial air samplers. These studies suggest that aerosol capture systems integrated with highly sensitive nucleic acid amplification for viral identification, when optimized for frequent air sampling and automated viral detection, can identify pathogen contamination in indoor air, sometime prior to the manifestation of symptoms [21].

Current airborne pathogen monitoring efforts deploy air collectors in remote and rural locations and require storage and transport of airborne collected samples to a laboratory facility for extraction, purification, and processing. Degradation of RNA on collected filter samples was cited as responsible for inconclusive detection results in some studies [20]. Storage of collected filter samples at low temperatures (−80^0^C) was required for preservation of samples when significant time was needed between collection and detection cycles [21]. These requirements substantially limit the scalability of these existing viral monitoring approaches, along with restricting their capacity for providing early warning for pathogen detection. Beyond filtration, air sampling of airborne viruses includes impaction, cyclone-based collectors, liquid impingers, condensation growth tubes, and electrostatic precipitators (ESPs), each with relative advantages and shortcomings [22, 40].

In this paper, we demonstrate a high flow rate electrostatic precipitator system capable of continuous operation integrated with miniature collection wells as counter electrodes for airborne viral collection. Reverse-transcriptase loop-mediated isothermal amplification (RT-LAMP) is then used for detection of captured viral matter within wells, due to its high accuracy, specificity, and simplicity of use [49, 50]. Small volumes of reagent (30 μ*L*) containing LAMP primers are delivered directly to the collection well sites, and on-chip heating of reaction wells is conducted using integrated planar heaters. A colorimetric change during the RT-LAMP amplification reaction is optically imaged to indicate the positive presence of viral matter (collected aerosolized model pathogen). This integrated detector demonstrates the viability of using the highly sensitive isothermal nucleic acid amplification (NAA) techniques, such as LAMP, for real-time detection of airborne pathogens, without the need for off-site processing currently widely used [20,21, 37]. Our initial prototype paves the way for future autonomous airborne viral detectors using NAA, noting that NAA is currently the gold standard for pathogen diagnostics at point-of care. We postulate that a large density of integrated microfabricated micro-volume collection wells can increase the capacity of the detector for long-term autonomous in-field operation.

## 2. Materials and Methods

### 2.1. Electrostatic Viral Aerosol Precipitation

While a variety of sampling techniques have been utilized towards collection of airborne viruses, approaches with high flow rates are preferred since the transfer larger volumes of air through the collection device, improves the possibility of pathogen detection [40]. Flow rates of 100 to 1000L/min have been suggested to be the practical range for the collection and detection of low concentration airborne biological agents [44]. Specific to air-liquid interfacing, Electrostatic Precipitation (ESP) based sampling has been demonstrated for airborne collection of viral laden aerosols into small volumes of liquid both for aerosol enrichment and to maintain virus viability [37, 40, 45, 46]. Collection efficiencies measured are shown to be inversely related to the ESP device flow rate, with a reported collection efficiency of 99.3 % at 1.2 L/min and 20% at 12.5 L/min, when tested with polystyrene particles 0.5 μ*m i*n diameter from one such system [46].

**Fig. 1** shows a high flow rate (100L/min) single-stage electrostatic precipitator (ESP) used in our device. Incoming aerosols are electrostatically charged and deposited within a set of grounded collection wells. In general, charging of aerosols utilizes the following two mechanisms: (1) field charging and (2) diffusion charging. Field Charging is governed by concentrations of ion bombardment and electric field strength due to applied corona voltage, while diffusion charging is dependent on the available number of ions and duration of exposure to the corona discharge field. Once charged, electrohydrodynamic transport of charged aerosol droplets causes their precipitation onto the grounded collection electrode surface [30]. While low collection efficiency is expected for such a high flow rate unoptimized collector, the on-chip integration of a highly sensitive detection technique (RT-LAMP) means that virions collected directly at the collection site can be detected at very low numbers (∼ 30 RNA copies) [42,49]. Longer sampling times are needed when airborne viral concentration is low.

**Figure 1:**
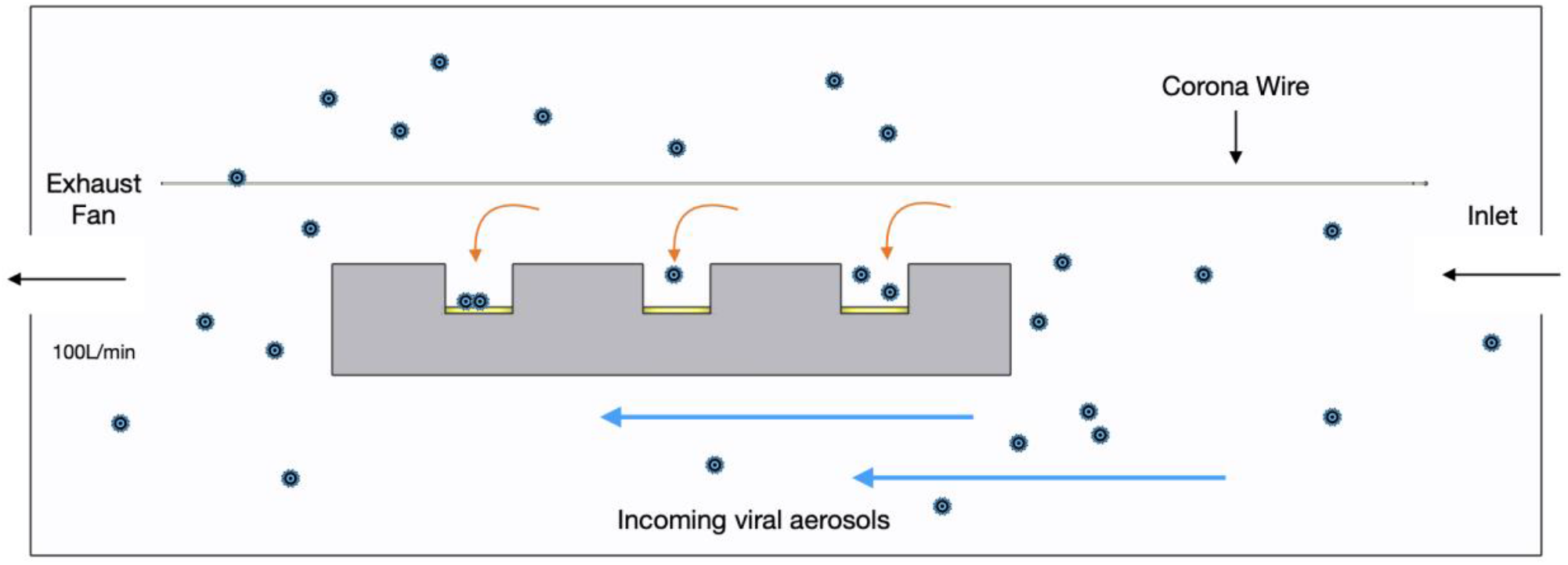
Cross-section diagram of the electrostatic precipitator.

### 2.2. ESP FEA Multiphysics Modeling

A finite element analysis (FEA) of the ESP was performed (COMSOL Multiphysics v. 5.2a) to verify the bioaerosol deposition on the well plate design. Laminar Flow Physics module was coupled with Corona Discharge Physics module to compute the fluid (air) velocity, space charge density, and electric field values used to solve for the size dependent charging of the incoming aerosolized particle stream. Next, Particle Tracing for Fluid Flow Module was used to verify particle collection on well plate surfaces [31,32]. The corona discharge model [32] solves for the transport of charge carriers through the domain using the Conservation of Charge Equation (Eq. 1) coupled with the Poisson Equation (Eq. 2).

The effects of drift and convection forces are related to the charge carrier transport equation as

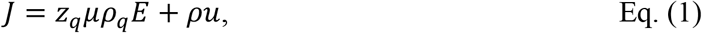

where *J* is current density (*A*/*m*^2^), *z*_*q*_is the charge number, μ *i*s the charge mobility (*m*^2^/*V. s*), *ρ*_*q*_is the space charge density (*C*/*m*^3^), *E i*s the electric field, *u i*s the fluid velocity (*m*/*s*), and

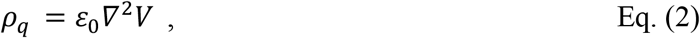

where, *V* is electric potential, *ε*_0_ is the permittivity of vacuum.

The transport equation terms (Eq. 3) are arranged from (Eq. 1-2) and describe the relationship between electric potential, space charge density and fluid flow. This domain equation does not contain information related to plasma generation or maintenance. All plasma physics is restricted to the boundary conditions at the electrode surface as

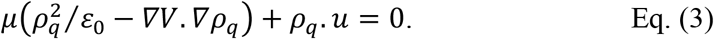

The generated plasma due to corona discharge results in a constant uniform electric field. Peek’s law (Eq. 4) [32] describes the onset of the electric field as a function of the radius of the corona wire as

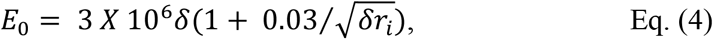

where *E*_0_ *i*s the breakdown electric field *V*/*m, δ i*s gas number density normalized to pressure of 760 torr and temperature of 293.15 K, *r*_*i*_ is the radius of the corona electrode.

The normal component of the electric field was provided as a first boundary condition at the surface of the corona electrode, where the applied voltage is 5 kV to the electrode component. A second boundary condition *V = 0 V* was provided at the collection surfaces. Additionally, no charges were considered at the inlet and outlet of the ESP model domain.

Next, to obtain the velocity and pressure profile of the flow field in the 3D domain, the air flow was modeled under incompressible flow conditions using the laminar flow module, which solves the Navier-Stokes’ equation for the conservation of momentum and mass. The inlet boundary condition was a laminar volumetric flow rate of 100 L/min, and the outlet boundary condition was set to atmospheric pressure. The wall boundary conditions were set to no-slip. Numerical particle tracing in fluid flow was used to investigate the behavior of particles inside the ESP. The particles were defined as spherical, with a mean diameter of 0.5 μm with the density of 1000kg/m3. The number of particles at the inlet was set to 10000.

The particle positions were solved using second-order equations of motion, following Newton’s Second Law affecting the particles, drag forces due to air flow, and the electrical forces. The drag force is described by the Cunningham-Millikan-Davis model (Eq. 5) as

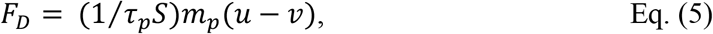

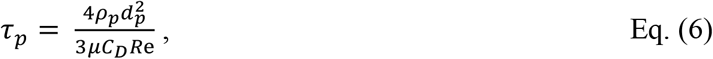

where *F*_*D*_ is the drag force; *τ*_*p*_ is the particle velocity-time response; *ρ*_*p*_ is the density of the particles, *d*_*p*_ is the particle diameter, *C*_*D*_ is the Cunningham correction factor, and *Re* is the Reynolds Number, and *S* is the drag correction factor defined as

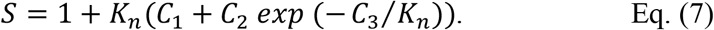

The electrical force (Eq. 8) due to accumulated charge on each particle is

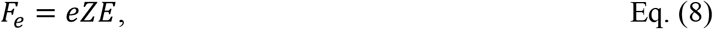

where *F*_*e*_ is the electrical force, *e* is the elementary charge and *Z* is the accumulated charge number for each particle. The particle charging rates are given by the Lawless model that includes the field and diffusion charging process (Eq. 9) as

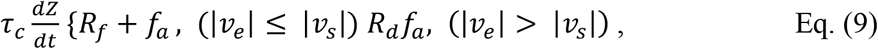

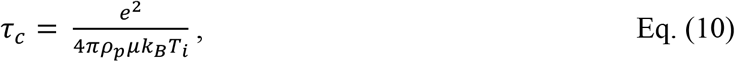

where *τ*_*c*_ is the charging time, *k*_*B*_ is Boltzmann’s constant, and *T*_*i*_ is the ion temperature; *R*_*f*_ and *R*_*d*_ are dimensionless values associated with the field and diffusion charging process, respectively, and *f*_*a*_ is an analytical fitting function that couples the two charging rates.

Finally, stationary studies were used to run the corona discharge, laminar flow, and charge transport models. Time dependent studies were used for particle tracing in fluid flow. **Fig. 2** presents the simulated particle capture across the well-plate collection surface. Images of the actual well-plated electrode designs are shown in **Fig. 2** inset.

**Figure 2:**
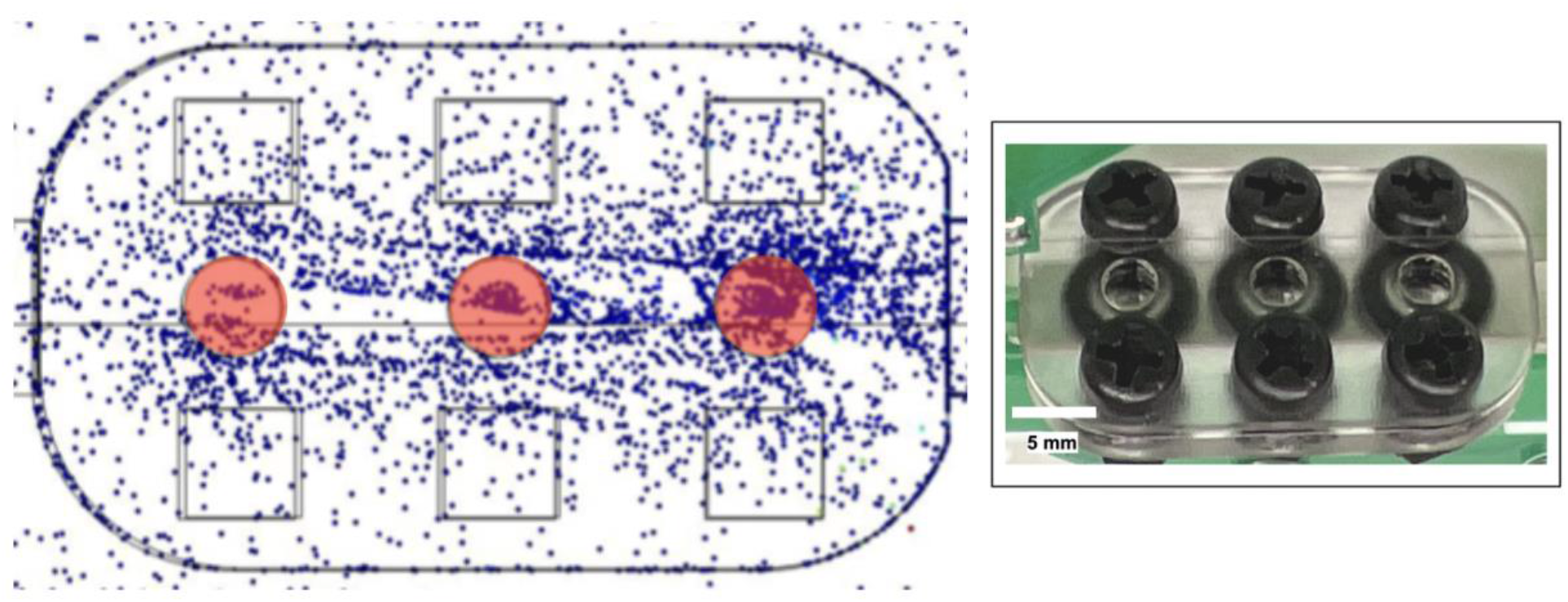
Electrostatic particle capture on well-plated collector (well area highlighted in red) in COMSOL – laminar volumetric flow rate is set to 100L/min and number of particles at the inlet are set to be 10,000.

### 2.3. Experimental Characterization of ESP

Evaluation and experimental characterization of the electrostatic precipitator collection efficiencies was carried out in an aerosol testing chamber as shown in **Fig. 3**. Mono-dispersed test aerosols, consisting of 0.5 μ*m* in diameter fluorescent polystyrene sphere latex (PSL) particles (Thermo Scientific Fluoro-Max), were aerosolized using a Collison nebulizer (CH Technologies) and introduced into the testing chamber. Compressed air (40 PSI) was routed through a HEPA filter (Gelman Lab - 12144) to enter the nebulizer containing PSLs suspended in deionized (DI) water. Aerosolized PSLs were passed through a diffusion dryer (TSI Diffusion Dryer 3062) and introduced into the testing chamber. Two fans placed in the chamber provided continuous air circulation to maintain uniform particle concentrations. An optical particle counter (TSI DustTrak II) was used to monitor the concentrations within the chamber during experimental trials. Additionally, air within the test chamber was sampled through a Mixed Cellulose Ester (MCE) filter (Millipore - 0.45 μm pore size) utilizing an external sampling pump (GilAir Plus) at 425 mL/min. ESP sampler collectors and filters within the chamber were operated for 15 min for each experimental trial. Three (3) experimental trials were conducted to characterize collection efficiencies for the well plate collector.

**Figure 3:**
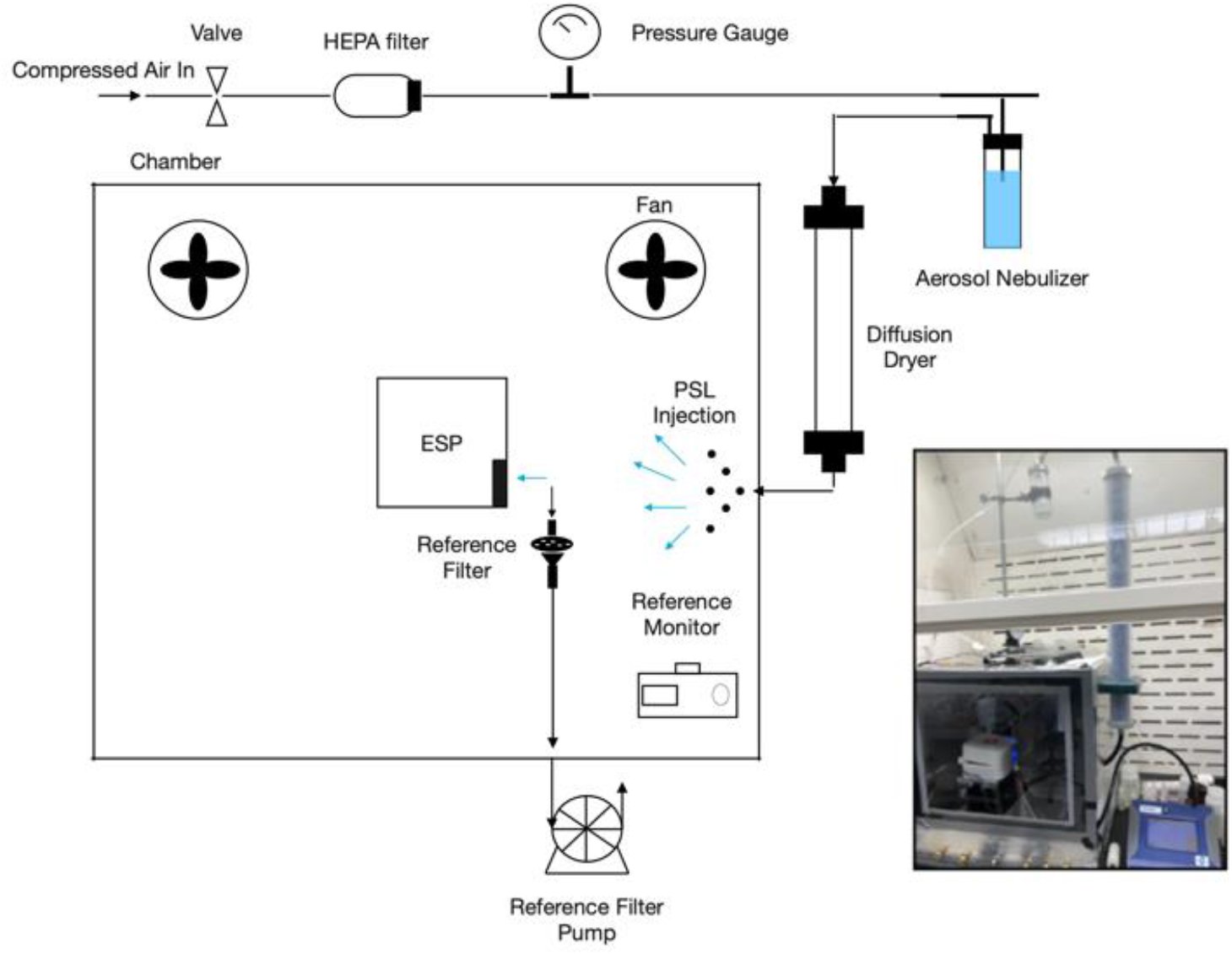
Diagram and picture (inset) of the aerosol testing chamber used for the experimental evaluation and calibration of the ESP sampler system collection efficiency.

PSLs captured on the grounded counter electrode collection surface within the ESP sampler, and the reference filters from each experimental trial, are imaged using an epi-fluorescence microscope (Zeiss Axio Scope A1). The resulting micrographs were quantified using algorithmic counting (Python OpenCV Library). For density analysis on the planar surface, five (5) images from each of the six uniformly distributed locations on both sides of the planar collection surface were quantified to determine count density at each location. For the reference filter, ten (10) images from five (5) locations distributed across the filter were similarly imaged. The reaction well masks were 3D printed using a resin printer (Stratasys Object30) and held over the grounded collector surface using O-rings and plastic screws, to create a removable well mask. For data analysis, fluorescent microscopy imaging and processing steps (**Section. 2.3.1**) were used to measure particle deposition efficiency within well sized area after removal of the reaction well masks.

#### 2.3.1. Image Processing and Collection Efficiency Calculation

A representative optical micrograph from fluorescent imaging is shown in **Fig. 4.a**. Pre-processing of raw image data started with a non-linear filtering process followed by thresholding of images using Otsu Technique [47, 48]. Post thresholding, binary images were subject to morphological operations to accurately identify particle boundaries and differentiate image pixels containing particles from background utilizing OpenCV libraries [33]. Contour diameters were measured for each individual particle feature along with frequency of occurrence. A histogram plot of particle contour diameters and corresponding frequencies for the representative image frame is shown in **Fig 4.b**. Contour sizes with low frequencies (particle agglomerates) were ignored. Finally, the frequency threshold was selected so that the median value of contour diameters for counted particles converges toward known PSL aerodynamic diameter of 0.5 μ*m*. As an example, for representative frame in **Fig. 4.a**, 2831 particles were counted, and median contour diameter was 0.513 μ*m*. This procedure is executed for all imaged optical micrographs to determine the total count and count density of PSL particles from image data collected on ESP collection surfaces, well-plate collector, and MCE reference filter surfaces.

**Figure 4:**
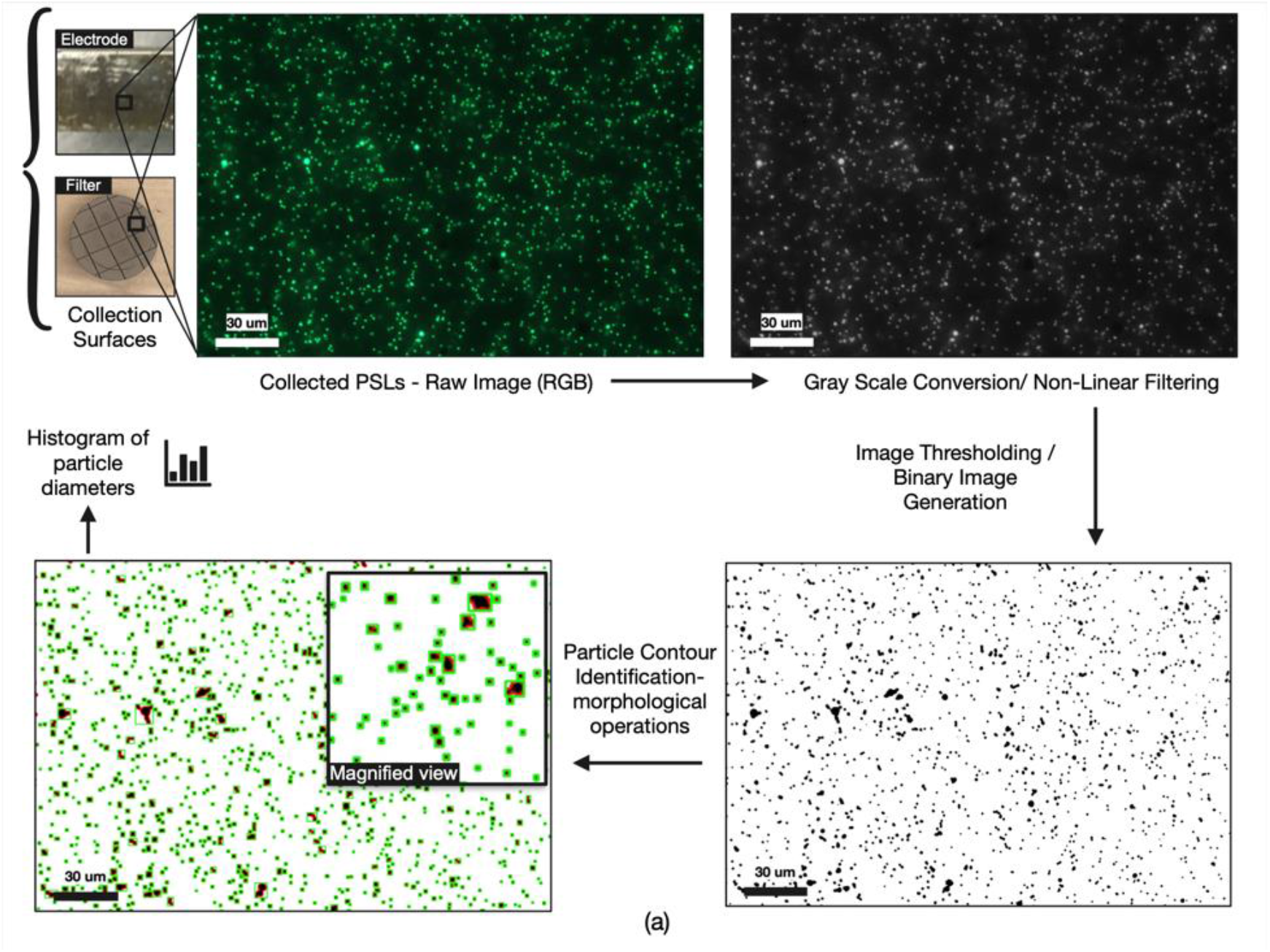
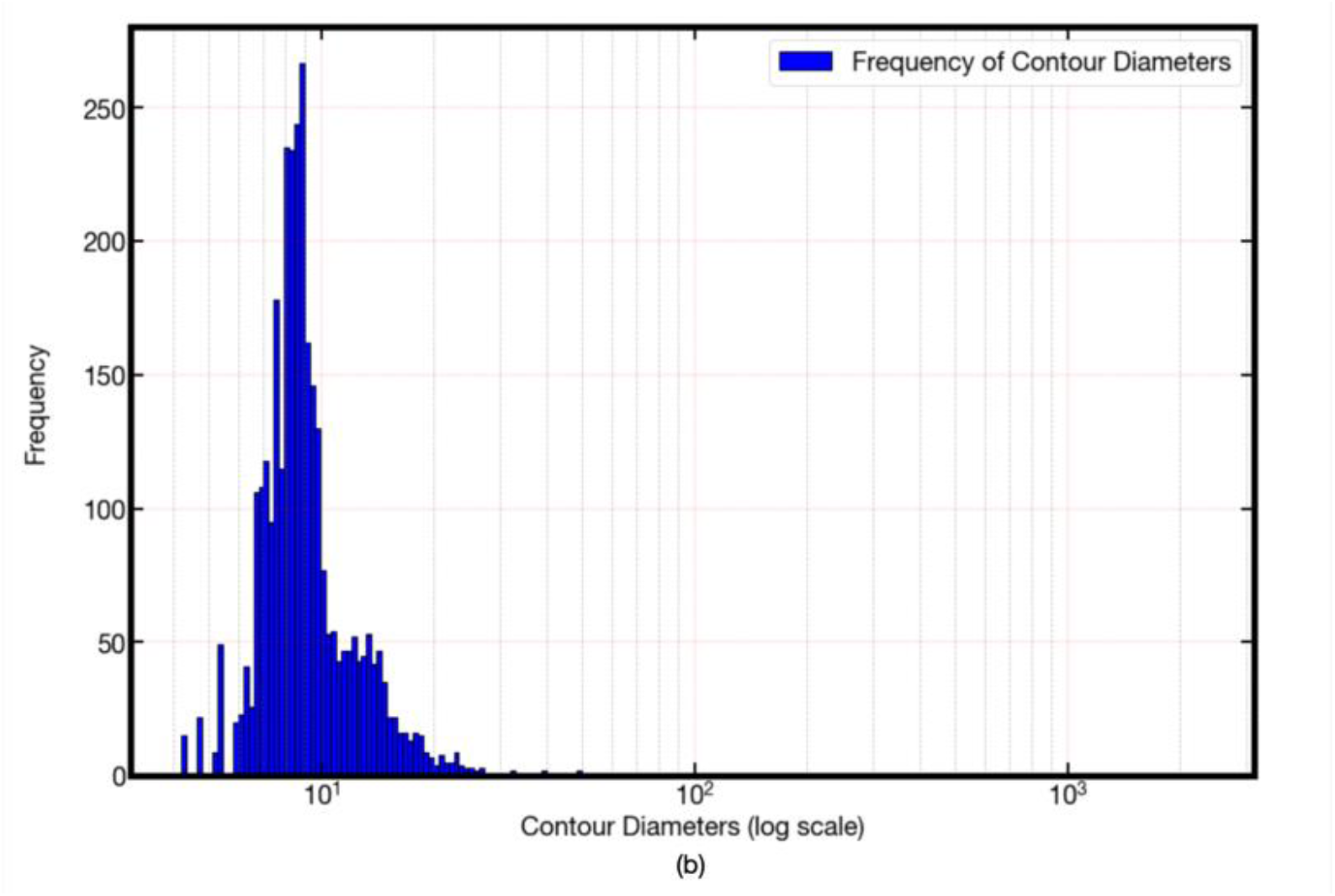
a) Optical Micrograph Image and image processing steps for particle counting. b) Histogram plot of particle contour diameters and corresponding frequencies for the representative image frame; In representative frame, **2831** particles counted, and median contour diameter was **0.513** μ*m*.

Total number of collected particles on collectors and filters is calculated by multiplying the count density averages with the surface area of electrostatic collectors, and filter surface areas, respectively. To determine the particle collection efficiency for the ESP sampler system, total number of collected particles on ESP collection surface and reference filters are normalized to account for differences in surface area and flow rates.

Collection Efficiency for the well-plate collector surface is calculated according to Eq. 11.

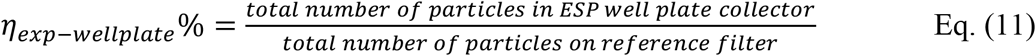

### 2.4. VLP Synthesis

As a first step, we generated VLP using the HIV vector pNL43r-e-as the packaging vector and an expression plasmid containing SARS-CoV-2 spike. The packaging vector contains the entire HIV genome with 2 mutations that result in nonfunctional envelope and rec proteins and thus the virus was considered noninfectious and VLP were produced under BSL2 conditions. One advantage of this packaging vector is that the resulting VLP contains a copy of the viral RNA of the HIV genome, which we will exploit as a marker to track aerosolized VLP by nucleic acid amplification. Addition of the second plasmid containing SARS-CoV-2 spike in the transfection results in the VLP containing an authentic version of the viral envelope protein within the VLP membrane. As such, these VLP spikes are almost identical in size (∼100 nm) and surface composition (spike and membrane lipids) to infectious SARS-CoV-2.

Plasmids containing SARS-CoV-2 spike and the packaging construct pNL4-3. Luc.R-E− were co-transfected into 293T cells, as previously described [34-36]. VLP were quantified by a LFA assay for HIV p24 (GoStix Plus, Takara). In this case the number of virions per volume is calculated by the approximate stoichiometry of p24 in VLP (∼2000/virion). Based on p24 concentration, the VLP solution concentration was estimated to be 3.2 X 10^9^ virions/mL in the PBS buffer.

#### 2.4.1. VLP Aerosolization and In-Situ Viral Detection using RT-LAMP

The aerosolization experiments were performed in a separate bio-aerosolization BSL2 chamber (see **Fig. 5**). Aerosolization of VLP utilized the Bio-Aerosol Nebulizing Generator (BANG) from CH Technologies. The pressurized air flow into the nebulizer was 4.72 L/min and the sampling pump flow rate for collection on the filter was 425 mL/min. A fan was placed in the BSL2 cabinet to provide continuous air circulation to maintain uniform aerosol concentrations. ESP collectors and filters within the chamber were operated for 30 min for each experimental trial.

**Figure 5:**
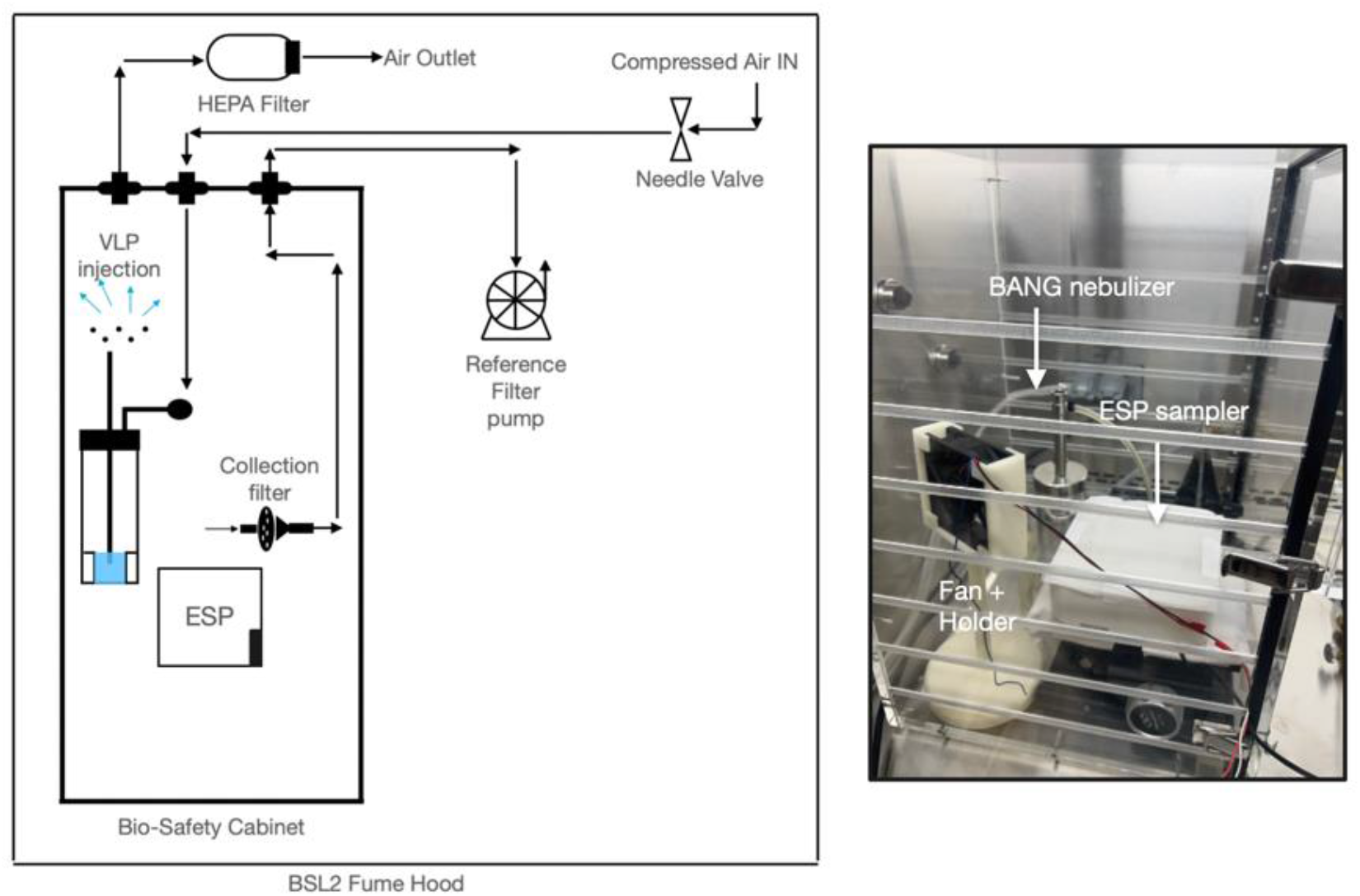
Diagram and picture (inset) of bio-aerosolization testing chamber used for used for experimental aerosolization and collection of VLPs.

The ESP collector surface for VLP deposition was modified to integrate two 50 *μL* collection wells (Thermo Fisher Scientific AB0620). The wells were metallized with gold using a sputter deposition system (CVC SC-4000 RF Magnetron Sputtering System) to provide connectivity to the ground terminal of the ESP device to shape the electric field for deposition into the well.

Experimental collection (3 trials) of aerosolized VLPs onto the well-plate collector surface and reference filters were conducted for 30 min in the BSL2 aerosolization chamber. The estimated concentration of VLPs in the chamber was 59,260 VLPs/L or 59 x 10^6^ VLPs/m^3^ of air (**Section. 2.4.4**). Following VLP deposition, a control well was fixed to the collector surface that served as a control well. LAMP reagents (30 μ*L*) required to detect VLPs were manually added to both the collector and control wells. As shown in **Fig. 6**, the collector plate assembly was heated using heat block (Fisherbrand Mini Dry Bath) to perform the isothermal RT-LAMP reaction at 65^0^C for 30 min directly at the collection site, verifying on-chip detection. While the colorimetric change is visible, the gold sputter layer reduces the transparency of the collection well. (**Section. 3.2**).

**Figure 6:**
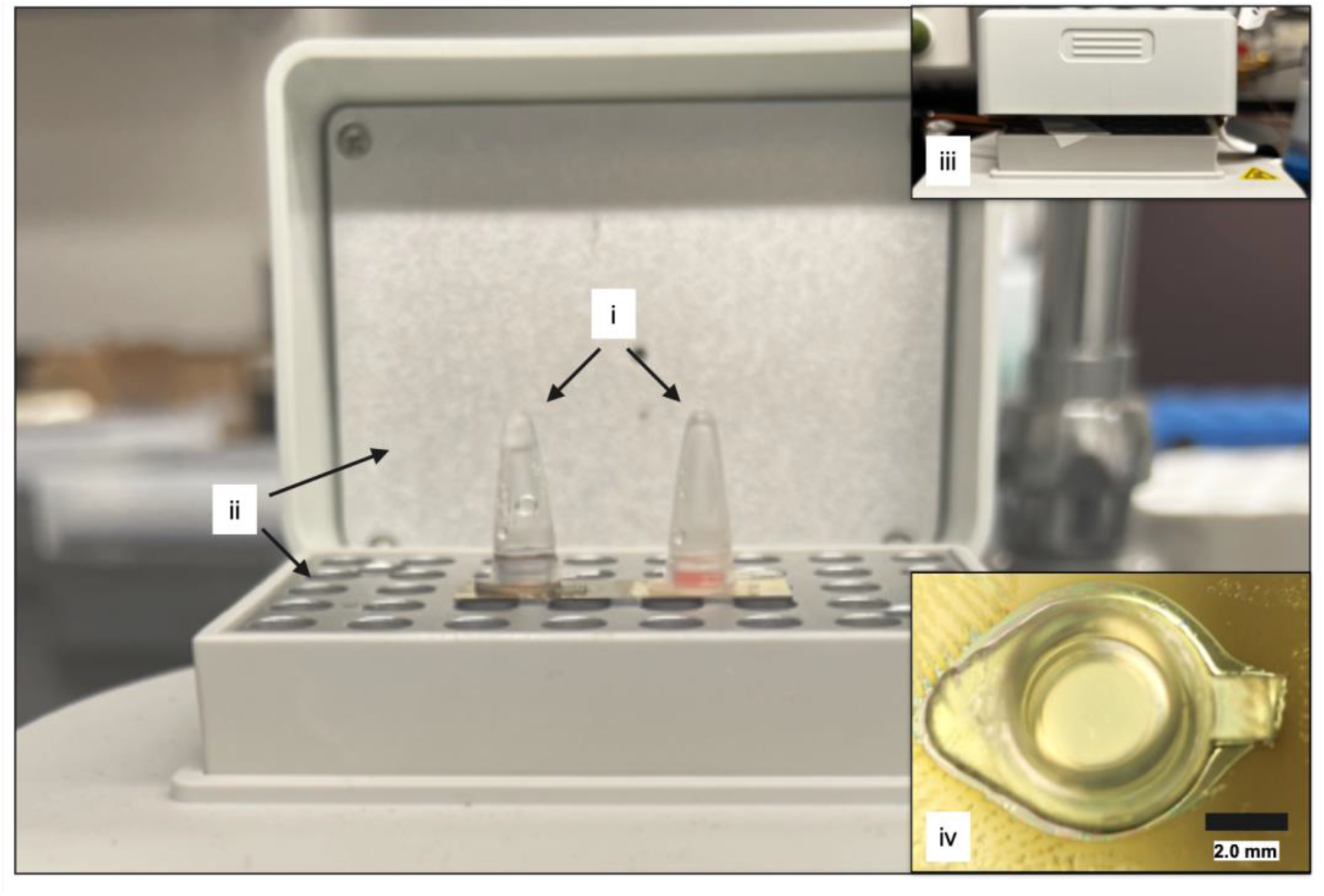
The collector plate assembly heating (Fisherbrand Hotplate) to perform the isothermal RT-LAMP reaction at 65^0^C for 30 min.: (i) Hermetic cap to prevent evaporation (ii), Heated surfaces, (iii) Hot plate closed during heating, and (iv) Metallized collection well – top view.

#### 2.4.2. Real-Time Detection of Viruses using RT-LAMP

We demonstrated the feasibility for on-chip RT-LAMP based detection of airborne captured viral RNA using a miniaturized enclosure for the collector well heating with *in situ* integrated heaters. The integrated heating/detection unit, (See **Fig. 7**), consists of a transparent, laser-cut acrylic enclosure surrounding the collection plate. The small volume of enclosed air facilitates uniform heating of collection wells, and the transparent setup ensures uninhibited optical readout of colorimetric change of LAMP reagents. Planar heating elements fabricated using printed circuit board (PCB) techniques (50 cm × 25 cm) are incorporated for independent, double sided heating of enclosure. The collection plate containing airborne captured virus is in contact with the bottom heater with a set point of 62°C, while the top heater, provides the convective heating of the ambient air surrounding for accelerated heat delivery to the reagent mixes contained in wells. The temperature on the collection plate surface is monitored using a J-type thermocouple (34307 – Keysight) cable and data acquisition unit (Keysight 34972A). The temperature feedback is provided through a DC voltage supply (TENMA 72-8690A/ Keysight E36318) to maintain temperature at 62^0^C for the reaction time of 30 min as shown in **Fig. 8**. HD Images (Cannon EOS Rebel T3) were captured every minute for the 30 min reaction time for real-time monitoring. Both wells were then metallized with gold using a sputter PVD deposition system (CVC SC-4000 RF Magnetron Sputtering System) to facilitate grounding and ESP deposition. Small pieces of aluminum foil were used to create a viewing window by shielding the front side of the collection well during gold deposition, for clear optical readout (**Fig. 7 (ix)**). Electrostatic capture of aerosolized VLPs were performed under BSL 2 conditions, identical to the aerosolization procedure and controls described above and was immediately followed RT-LAMP reaction. Quantification of optical readout is described (**Section. 2.4.3**) and utilized for both detection in real-time and at the reaction end point (**Section. 3.2**).

**Figure 7:**
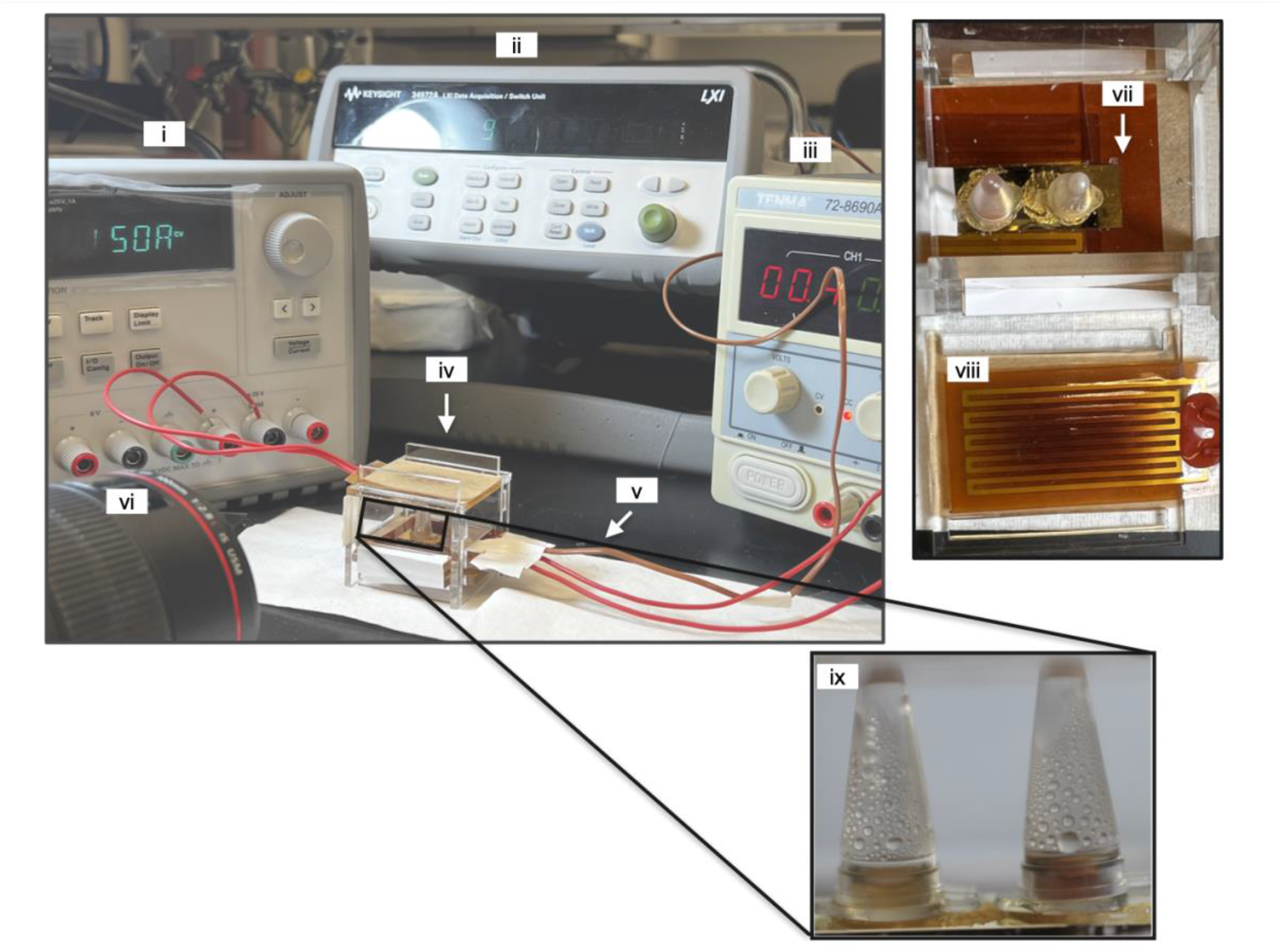
On-chip heating and RT-LAMP detection unit: (i) DC voltage source – top heater control, (ii) Temperature readout, (iii) DC voltage source – bottom heater control, (iv) Reaction enclosure, (v) Thermocouple connections, (vi) CCD camera, (vii) Bottom heater, (viii) Top heater, and (ix) Capped reaction wells – front view.

**Figure 8:**
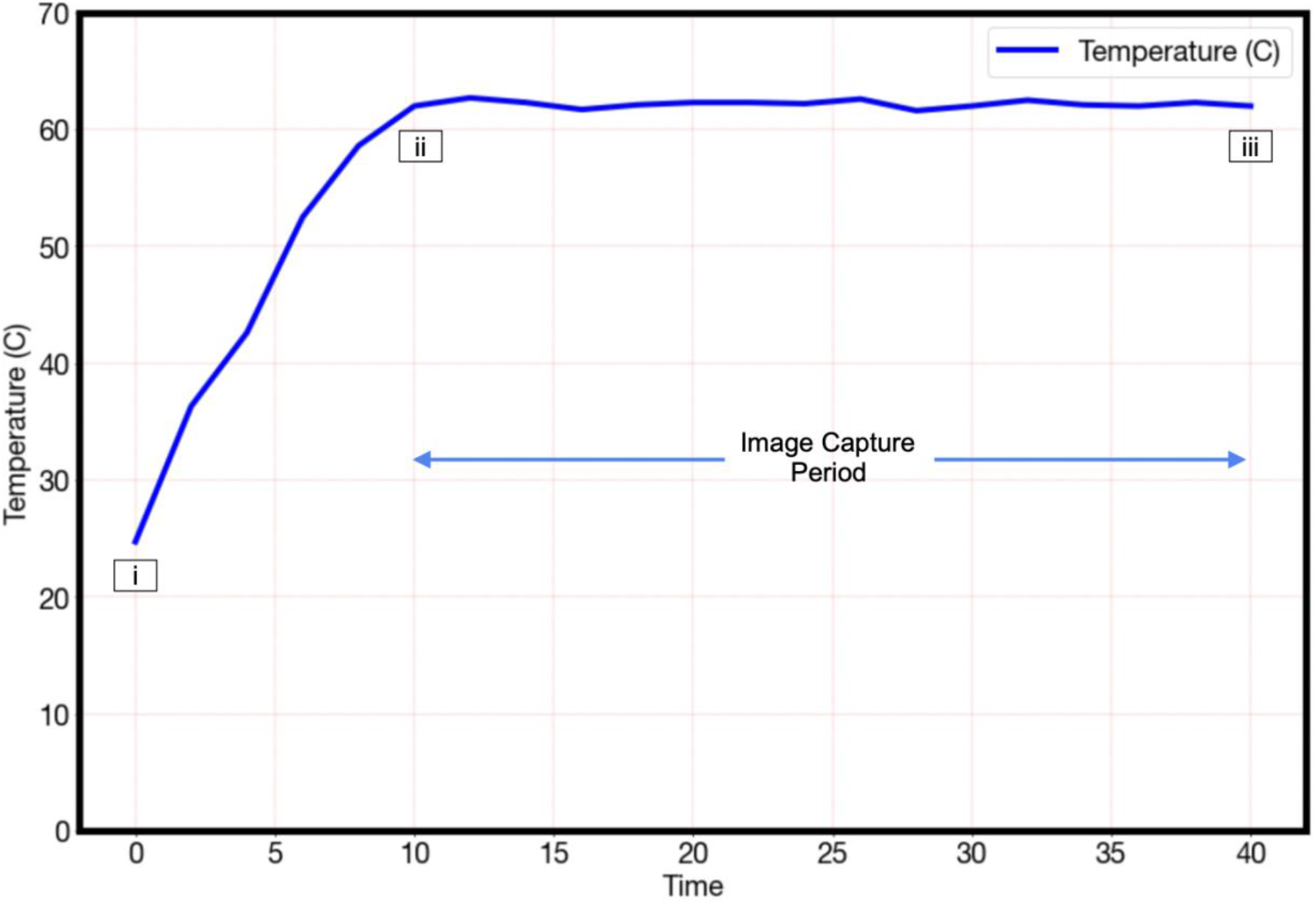
Closed-loop temperature control during RT-LAMP reaction using planar heaters (temperature set point: 62^0^ C).: (i) Start of the pre-reaction warmup, (ii) Warmup complete, consider start of RT-LAMP reaction, (iii) End of RT-LAMP reaction.

#### 2.4.3. Image Processing for Colorimetric Readout

Representative images of colorimetric readout from RT-LAMP reaction of airborne collected virus like particles, along with subsequent processing steps are presented in **Fig. 9**. The detection procedure was executed in Python utilizing OpenCV packages for image data extraction, analysis, and plotting. First, regions of interest were selected to avoid imaging artifacts and highlight regions of uniform color signal, both in control and collection of well image data sets. A histogram of green (RGB) channel was extracted, and peak response values are identified in each region of interest, for both control and positive test reaction image data. Green channel peak responses for end-point detection of airborne VLP capture using RT-LAMP, were used as a relative change before and after heating, for both positive test collection well (T) and control wells (C). For real-time detection of colorimetric change, images were acquired at 1 min increments for the 30 min run-time of the RT-LAMP reaction.

**Figure 9:**
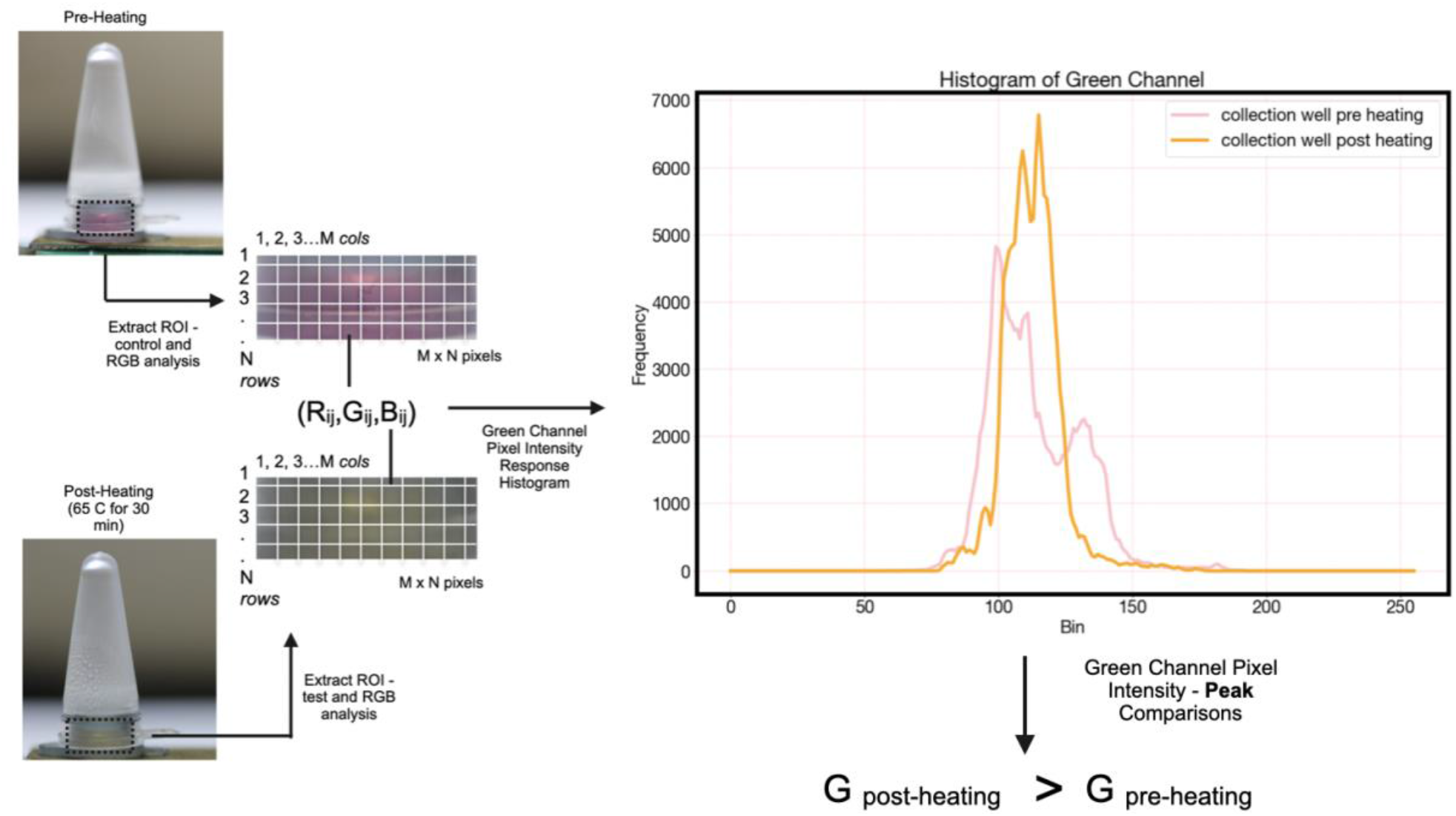
Steps adopted for algorithmic processing and quantification of colorimetric readout during RT-LAMP – for real-time and endpoint detection of airborne captured viral RNA.

## 3. Results

### 3.1. Experimental ESP Characterization

As detailed in **Section 2.3**, fluorescent polystyrene beads (0.5 μ*m*) are utilized to characterize the performance of our ESP device. For reference, we also present the results for the collection efficiency corresponding with the maximum electrode surface area permissible within our ESP i.e., a double-sided collector 30 mm × 5 mm. Experimental Collection Efficiency values for well and double-sided collector are calculated and summarized in **Table 1**.

**Table 1:** Experimental ESP collection efficiency values – individual wells and double-sided collector with corresponding collection surface areas.

| *Collector Electrode* | *Collection Efficiency (%)* | *Collection Surface Area ($\text{mm}^2$)* |
| --- | --- | --- |
| *Well 1 (L)* | **0.131** | **7.06** |
| *Well 2 (M)* | **0.143** | **7.06** |
| *Well 3 (R)* | **0.168** | **7.06** |
| *Double Sided Collector* | **0.59** | **300** |

Although our results show low collection efficiency (approximately 0.15 %) into the integrated reaction wells, we note again that low collection efficiencies are common at high flow-rate ESPs, and at our flow-rate of 100 L/min the system is still expected to precipitate 1-2 virions/min at an ambient concentration of 10 virions/L (e.g., see [51] for relevant concentrations).

### 3.2. Experimental Characterization of VLP Aerosolization

As detailed in [42] we developed a BSL2 model for airborne viruses using VLP containing the SARS-CoV-2 spike protein in the membrane. In this model, the VLP contains a copy of the HIV genome in the form of RNA, which includes the HIV pol gene. A RT-LAMP assay has been designed for the detection of these aerosolized VLPs. In this assay, the color changes from pink to yellow in the presence of RNA/DNA amplification due to a decrease in pH of the reagent solution. To determine the concentration of aerosolized VLPs in the BSL2 chamber, VLPs were extracted from the filter surface in an extraction buffer of 200 μ*L*, as shown in **Fig. 10**. This stock concentration was verified using the ELISA assay. An aliquot of 1μ*L w*as taken from the extraction buffer containing a fraction of the total collected VLPs and then amplified using RT-LAMP assay to detect the presence of VLP RNA. A procedure of successive dilutions and assay detections was performed until a negative colorimetric readout was reached from the RT-LAMP assay. Agarose Gel Electrophoresis was conducted at each step, to confirm RT-LAMP amplification when VLP are present. Next, concentration in the chamber was calculated based on number of aerosolized VLP on the full MCE filter (0.45 μM) surface area. Using this procedure, we calculated that the total number of VLP collected by the filter, assuming 100% extraction, was approximately 800,000. Accounting for reference filter sampling rate of 0.425L/min and operating time of 30 min, concentration in chamber is calculated to 59,260 VLPs/L or 59 × 10^6^ VLPs/m^3^ of air.

**Figure 10:**
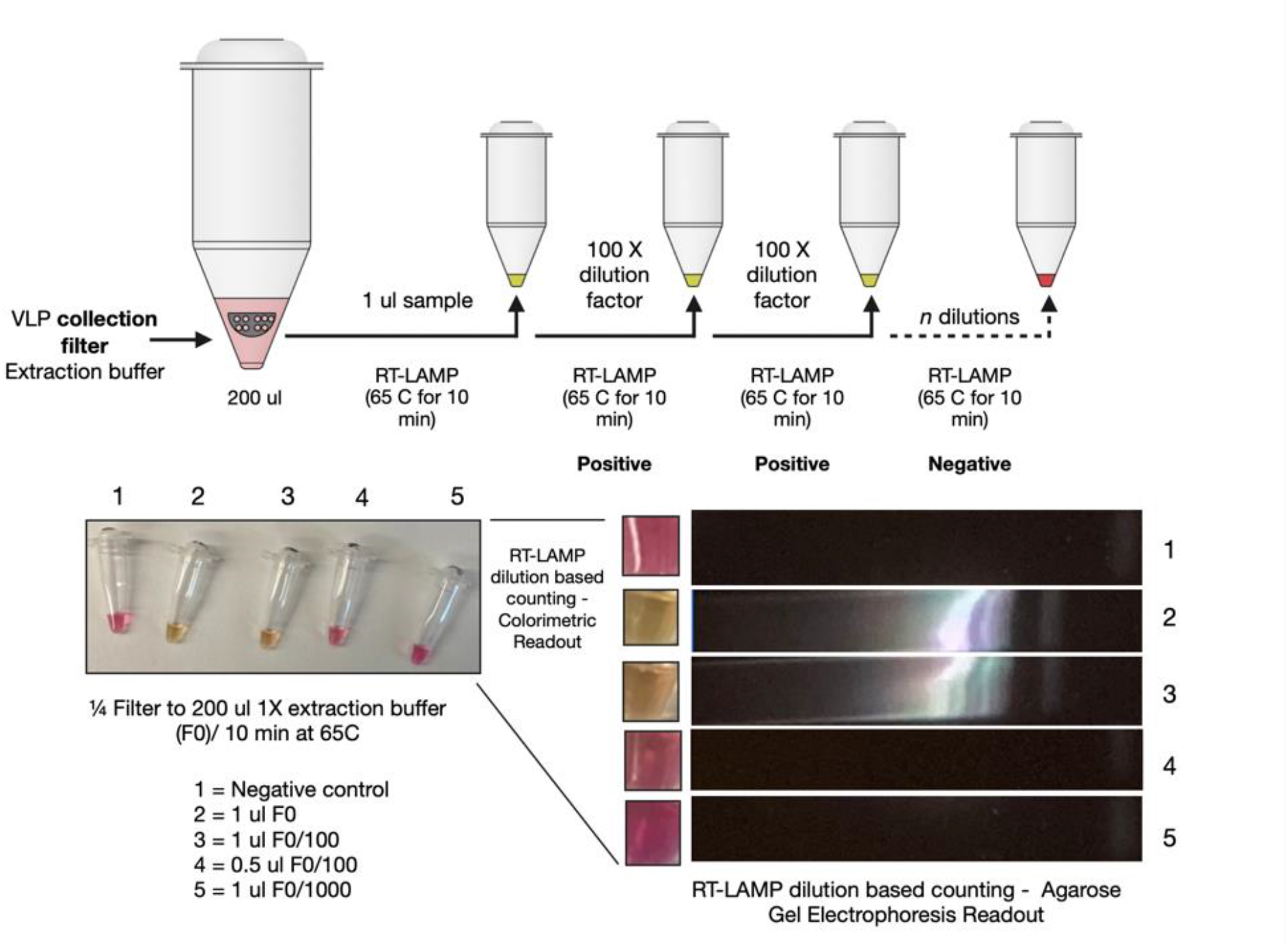
a) Colorimetric RT-LAMP assay; b) SYBR green stained agarose gel of the RT-LAMP reaction shows the presence of nucleic acid amplification when VLPs are present.

### 3.3. Electrostatic Precipitation and Real-time Detection of VLPs

The model pathogen (VLPs) was aerosolized and deposited inside the reaction wells as described earlier. **Fig. 11.b** shows the colorimetric RT-LAMP assay conducted *in-situ*. The results show a representative response from the detected VLP RNA, and a corresponding control well. The difference of contrast between the control well and the deposition well is due to the absence of a counter-electrode metal coating on the control well.

**Figure 11:**
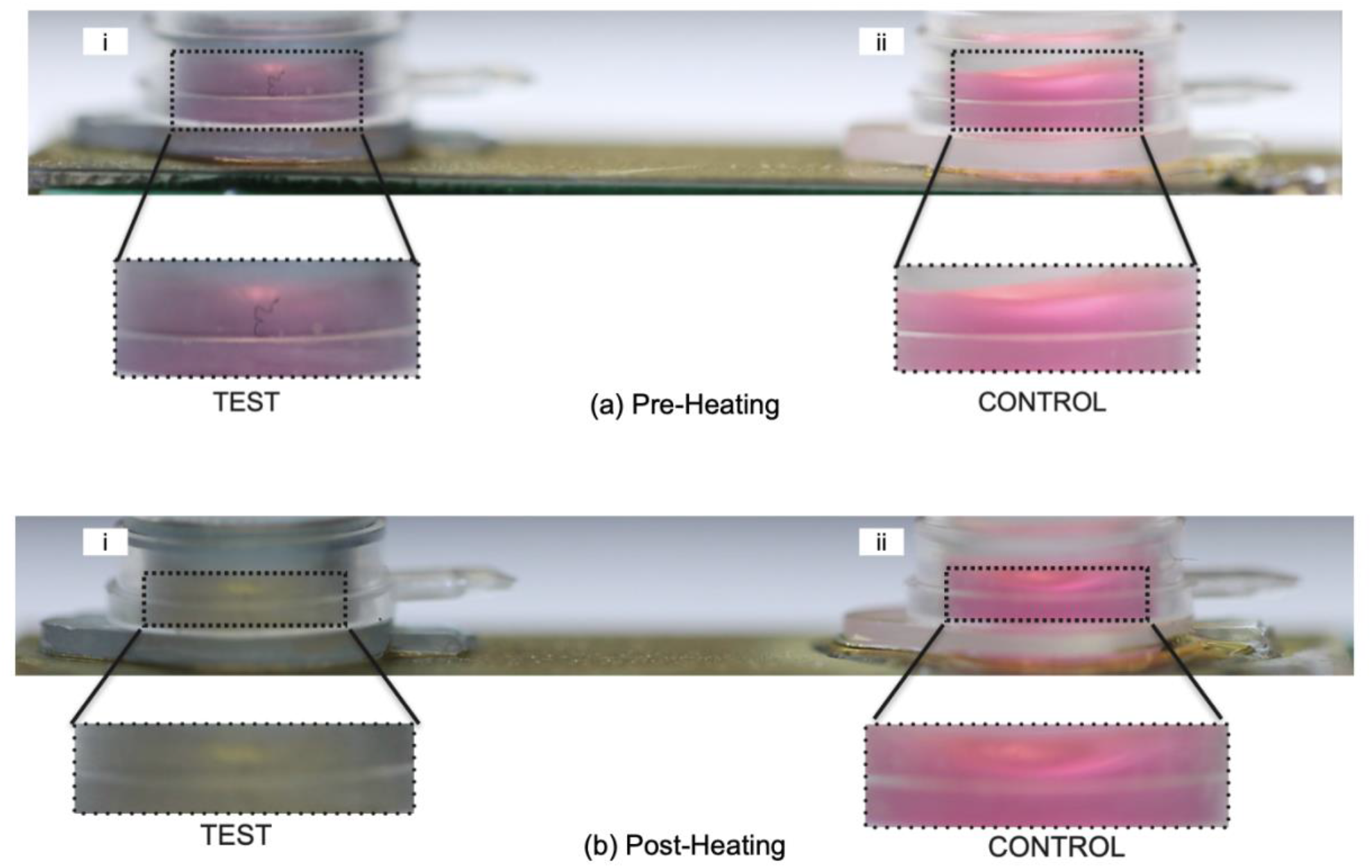
a) Colorimetric RT-LAMP reaction on VLP collector well – pre-heating; b) Colorimetric RT-LAMP reaction on VLP chip collector – post heating (65^0^C for 30 min.): (i) Collection well, and (ii) Control well.

Relative change in green channel colorimetric readout for the positive test reaction (T) and control reaction (C) wells using lab-grade (Fisherbrand Mini Dry Bath) hot plate are shown in **Fig. 12.a** as quantification of end-point detection of captured VLPs. The real-time colorimetric green-channel response data collected during on-chip detection using *in-situ* heater/collection unit is shown in **Fig. 12.b**.

**Figure 12:**
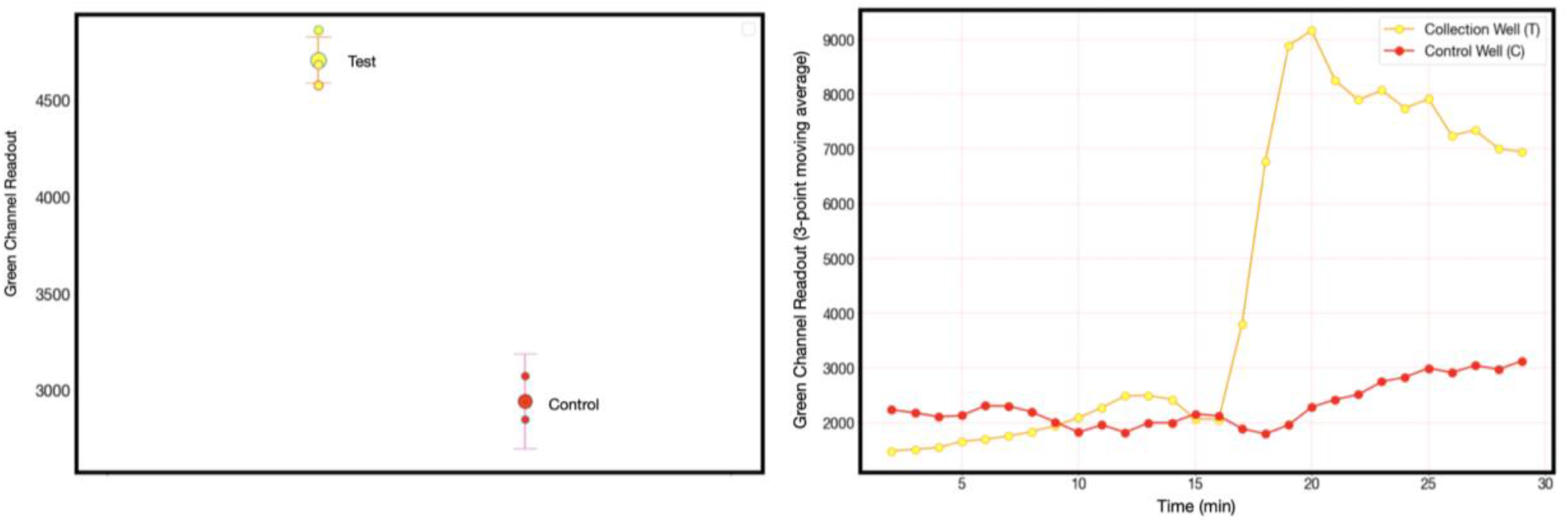
(a) Relative change in colorimetric readout for the positive test reaction (T) and control reaction (C) - quantification of on-chip end-point detection of captured VLPs; (b) Graph from a representative real-time colorimetric output RT-LAMP in the reaction well with precipitated VLPs during LAMP reaction.

## 4. Discussion

The integrated electrostatic collection well-plate and miniaturized heating/detection unit detailed in this paper demonstrates the capacity for airborne capture and *in situ* viral detection in real-time (< 30 min). This multi-well integrated collection/detection platform verifies viral detection *at the collection site*. We refer to this as *one-pot* NAA detection. We want to underscore that our experiments show the robustness of the one-pot reaction method using the developed LAMP assay for viral aerosol detection. We realize that the ESP capture system was designed to trade efficiency for high flowrates and leaves room for improvement, while the heating reaction well was implemented using simplified feedback temperature control. Because we are conducting the NAA reaction directly in the collection well, we can circumvent the usual extraction losses from viral extraction off the collection surface using an extraction buffer and the subsequent concentration of viral RNA within the extract solution. We anticipate therefore that the number of viral copies per well for the one-pot reaction is close to the sensitivity of the RT-LAMP assay i.e., less than 30 RNA copies [42,49], a result we have confirmed experimentally using stock VLP solution. This will be further analyzed using aerosolized VLPs in (ongoing) future work.

## 5. Conclusion

In this paper, we have demonstrated a high flow rate of electrostatic sampler with an integrated multi-well heating and detection system for the near real-time (< 30 min) detection of airborne pathogens using an aerosolized model for viral pathogens (VLPs). Direct electrostatic air capture of VLPs followed by *in-situ* LAMP based detection of viral RNA was demonstrated without the need for RNA extraction and purification. The low volume of reagent (30 μ*L*) required from using a miniaturized well collector) ensures a low cost-per-reaction for each detection cycle. Taken together, our high flow rate collection and detection scheme demonstrates a promising novel system architecture for a scalable bio-surveillance sensing system that is applicable to monitor indoor spaces for viral pathogens. Along with further miniaturization steps, fully automatic system operation is the scope of future work.

## Abbreviations

FEM: Finite Element Modeling
RT-LAMP: Reverse Transcriptase Loop Mediated Isothermal Amplification
ESP: Electrostatic Precipitator
NAA: Nucleic Acid Amplification.

## Acknowledgements

This work was partially supported by a grant from the UIC Center for Clinical and Translational Sciences (CCTS) to MC and IP. The work was also partially supported by contract 200-2016-91153 to IP from U.S. Center for Disease Control and Prevention/NIOSH. The findings and conclusions in this manuscript are those of the authors and do not necessarily represent the views of CDC/NIOSH. The mention of company names or products does not constitute endorsement by CDC/NIOSH.

## Notes

### Competing Interest Statement

The authors have declared no competing interest.

